# The N-terminal PAS domain directly regulates hERG channel gating in excised, inside-out patch-clamp fluorometry (PCF) recordings

**DOI:** 10.1101/2024.07.21.604498

**Authors:** Matthew C. Trudeau

## Abstract

Human ERG is a voltage-activated, K-selective channel whose physiological role is to drive action potential repolarization in cardiac myocytes. To carry out its role in the heart, hERG has specialized gating (opening and closing) transitions that are regulated by the internal N-terminal PAS and C-terminal CNBH domains. The PAS and CNBHD domains interact directly and this interaction is required for the characteristic slow deactivation (closing) of hERG channels. But it is unclear whether PAS remains globally attached or dislodges from the CNBHD during gating. Interestingly the direct PAS-CNBHD interaction can be formed *in trans* by co-expression of the PAS domain and hERG channels with a deleted PAS domain (hERG ΔPAS) in which the PAS domain is not attached to the channel with a peptide bond. *In trans* expression allows us to probe the biophysical mechanism for PAS domain attachment to the rest of the channel and in a broader sense allows us to test the mechanism for intracellular domain function in an ion channel, and test whether the PAS domain detaches or remains attached to the channel during gating. We report here that in excised patches from cells containing the hERG PAS domain fused to CFP and hERG ΔPAS channels fused to Citrine that 1) regulation of deactivation (slow deactivation conveyed by the PAS domain) was similar in on-cell and excised, inside-out patch configurations, 2) that regulation of deactivation persists for the lifetime of the patch (up to 30 minutes) in excised, inside-out mode, 3) that channel activity measured by activation of the channel with voltage pulses did not alter channel deactivation and 4) dual fluorescence and ionic current measurements using patch-clamp fluorometry (PCF) showed that only membrane patches containing PAS-CFP + hERG ΔPAS-Citrine had CFP and Citrine fluorescence and slow (regulated) deactivation, whereas control patches with hERG ΔPAS -Citrine had fast (unregulated) deactivation and Citrine fluorescence (but not CFP fluorescence) and control patches from hERG PAS-CFP - injected cells had neither currents nor CFP or Citrine fluorescence. Moreover, in PCF mode, we detected FRET from PAS-CFP + hERG ΔPAS-Citrine channels. Taken together, these results suggested that PAS - CFP remained associated with hERG ΔPAS-Citrine channels after membrane excision. We interpret these results to mean that the PAS domain was not dislodged from the channel despite mechanical (excised patch) and conformational (voltage) challenges and suggests that the PAS domain remained firmly attached to the hERG channel during gating.

## INTRODUCTION

The physiological role of hERG K channels is best understood in the heart, where they encode a repolarizing current that shapes and repolarizes cardiac action potentials (Sanguinetti et al, 1995; Trudeau et al, 1995). The importance of hERG and repolarizing cardiac currents is emphasized by familial mutations in the hERG gene that cause the Long QT type 2 cardiac arrhythmia syndrome that can degenerate into sudden cardiac death (Curran et al, 1995). hERG is similar to voltage-activated K channels in that it contains 6 transmembrane domains include the VSD and S4 helix and a pore domain. Like the CNG and HCN channels but unlike other K channels, hERG contains a CNBHD (Warmke & Ganetzky, 1994) and C-linker domain in its C-terminal region. Distinct from other K channels, hERG has an N-terminal Per-Arnt-Sim (PAS) domain (Morais Cabral et al, 1998) in the N-terminal region (Codding & Trudeau, 2023).

The PAS domain in hERG robustly regulates channel gating. The primary effect of PAS is to regulate and slow channel closing (or deactivation). Deletions of the PAS domain speed up channel gating by 5-10 fold, indicating loss of the regulatory effect (Morais Cabral et al, 1998; Spector et al, 1996; Wang et al, 2000; Wang et al, 1998). Point mutations within PAS speed up deactivation as if the PAS domain function were disrupted by the mutation (Morais Cabral et al, 1998). Missense mutations in PAS that are associated with LQTS also speed deactivation, indicating that hERG and the proper functioning of PAS is important for wild-type physiology (Chen et al, 1999; Gianulis & Trudeau, 2011). Like deletion of the PAS, mutations (Al-Owais et al, 2009) or deletion of the CNBHD speeds deactivation (Gianulis et al, 2013; Gustina & Trudeau, 2011). Deletion of both the PAS and CNBHD do not have an additive effect on deactivation, indicating they work by a convergent mechanism (Gianulis et al, 2013; Gustina & Trudeau, 2011) and indeed PAS makes a direct, intersubunit interaction with the CNBHD as measured with electrophysiology, biochemistry pull-down interactions, FRET hybridization assays (Gianulis et al, 2013; Gustina & Trudeau, 2011) and X-ray crystallography in EAG channels (Haitin et al, 2013) and CryoEM of a mostly full-length hERG channel (Wang & MacKinnon, 2017). Functional electrophysiology experiments showed that the PAS-CNBHD interaction was intersubunit (Gianulis et al, 2013; Gustina & Trudeau, 2011) and this was also shown in the CryoEM structure of hERG (Wang & MacKinnon, 2017) and EAG (Whicher & MacKinnon, 2016).

While the PAS interaction with the channel and CNBHD is consistent with being permanent (Gianulis et al, 2013; Gianulis & Trudeau, 2011; Gustina & Trudeau, 2009; Gustina & Trudeau, 2011; Gustina & Trudeau, 2013; Haitin et al, 2013; Morais Cabral et al, 1998) and undergoing small, Angstrom sized motions during gating in hERG as measured with cross-linking of non-canonical amino acids incorporated into the PAS domain or Molecular Dynamics (Codding & Trudeau, 2024; Stevens-Sostre et al, 2024; Trudeau, 2024) and transition metal FRET in ELK channels (Dai et al, 2018; Dai & Zagotta, 2017), (see Fig. 1, Model I) there has been speculation that the PAS domain mechanism involves the global PAS physically detaching and reattaching to the CNBHD during gating (Fig. 1, Model II) because hERG gating was modified by antibodies that bind to PAS (Harley et al, 2021). However, the distinction between the models has not been rigorously tested and here we perform experiments to distinguish between the two models.

**Figure 1.**
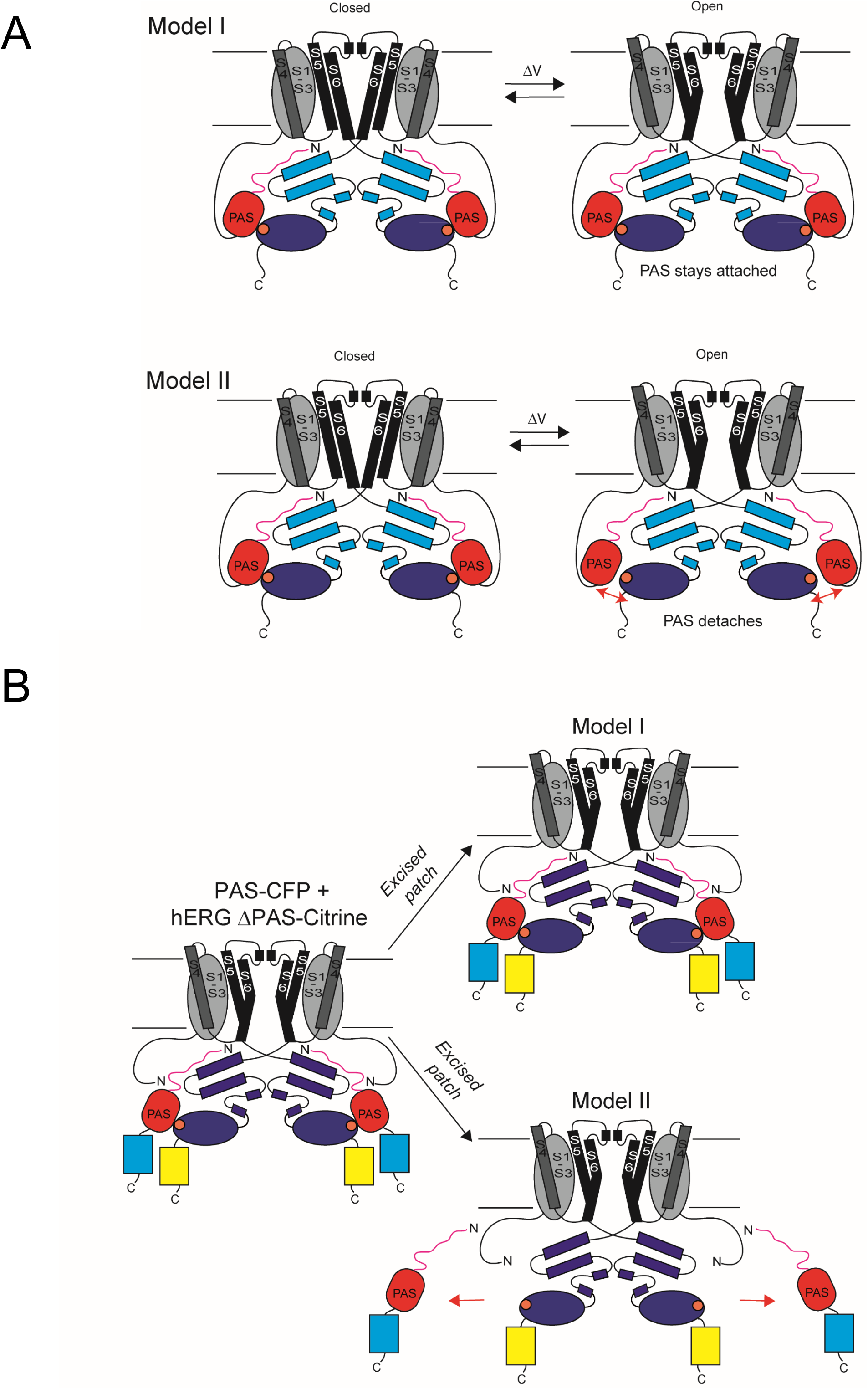
Scheme of PAS domain gating models and PAS *in trans* experimental design. A) Schematic for models of PAS domain gating. In Model I, the PAS remains globally attached during gating. This model allows for rotations or movements of PAS. In Model II, the alternative model, PAS disassociates and reassociates from the rest of the channel during gating. B) hERG channels formed by co-expression of PAS-CFP + hERG ΔPAS-Citrine. In Model I we anticipate that PAS-CFP would remain attached after patch excision into inside-out mode. In Model II we anticipate that PAS-CFP would detach, since it is not connected with a peptide bond to the channel, and then be diluted into the bath of the recording dish, leaving just hERG ΔPAS-Citrine channels. In excised, inside-out patches, channel deactivation in Model I is expected to be approximately 5-fold slower than in Model II, which can be used to distinguish between the models.

A powerful assay for measuring PAS regulation of the channel is the PAS *in trans* assay (Fig. 1B). In this approach, the PAS domain fused to CFP (PAS-CFP) was co-expressed with hERG channels lacking the PAS domain fused to Citrine (hERG ΔPAS -Citrine) as two separate mRNAs or cDNAs (Gianulis et al, 2013; Gustina & Trudeau, 2009; Gustina & Trudeau, 2011; Gustina & Trudeau, 2012; Gustina & Trudeau, 2013). The result of the *in trans* co-expression of PAS-CFP + hERG ΔPAS - Citrine is that the PAS domain protein interacts directly with hERG ΔPAS channels as measured by PAS regulation of channel deactivation and inactivation measured with electrophysiology and structural interactions as measured with biochemistry and fluorescence spectroscopy (Codding et al, 2020; Codding & Trudeau, 2019; Gianulis et al, 2013; Gianulis & Trudeau, 2011; Gustina & Trudeau, 2011; Gustina & Trudeau, 2013; Trudeau et al, 2011). This means that PAS does not require a peptide bond to the channel in order to exert its regulatory effect on gating and instead the PAS regulatory effect is due to its direct interaction with other channel domains (Gustina & Trudeau, 2009; Morais Cabral et al, 1998).

Here we test between a Model I and Model II mechanism for PAS regulation (Fig. 1A) by leveraging the PAS *in trans* experimental design (Fig. 1B). We propose that if PAS remained stably attached to the channel during gating then in excised, inside-out patches, PAS regulation would persist, i.e. deactivation would be slow, similar to that in wild-type hERG, after patch excision, whereas, in contrast, if PAS detached during gating or mechanical disruption of the membrane then PAS regulation would be lost, i.e. deactivation will become fast, similar to that in hERG ΔPAS channels.

Using the PAS *in trans* design, we found that deactivation remained slow and similar to that in wild-type hERG in the excised, inside-out patch-clamp configuration and furthermore, deactivation remained slow for the lifetime of the patch (at least 1/2 an hour) and in the face of a series of repeated voltage pulses that activated hERG. We next used patch-clamp fluorometry (PCF) to simultaneously measure function with ionic currents and structure with fluorescence (Dai et al, 2019; Dai et al, 2018; Dai & Zagotta, 2017; Trudeau & Zagotta, 2004; Zheng & Zagotta, 2000; Zheng & Zagotta, 2003). We found that in membrane patches from cells expressing PAS-CFP *+* hERG ΔPAS-Citrine we measured robust CFP fluorescence that was correlated with slow deactivation. Moreover, in PCF mode we measured FRET from membrane patches with PAS-CFP and hERG ΔPAS-Citrine, which also indicated that the PAS-CFP was associated with hERG ΔPAS-Citrine channels in excised patches. Together, our results suggested that the PAS domain remained associated with the hERG channel in excised, inside-out patches, which we interpret to mean that the PAS domain remained directly associated with the channel during gating, supporting the Model I view of gating in hERG.

## METHODS and MATERIALS

### Molecular Biology

hERG-CFP, hERG-Citrine, hERG ΔPAS-Citrine were all fused to the fluorescent protein at the C-terminus. PAS was fused to CFP after amino acid 135 as described in previous work (Gustina & Trudeau, 2009). Each channel contained the S620T mutant which ameliorates inactivation (Herzberg et al, 1998) for ease of analysis and to obtain robust currents. mRNAs were transcribed using the T7 mESSAGE mACHINE kit. A total volume of 49 nl of mRNA injected with a microinjector (Nanoject). Oocytes were surgically harvested from *Xenopus* (Nasco or NXR) and incubated in ND-96 solution at 16 degrees Celsius for up to 5 days. For PAS CFP + hERG ΔPAS, mRNAs were injected at a ratio of 2:1, as in previous studies (Gustina & Trudeau, 2009; Gustina & Trudeau, 2011).

### Electrophysiology and Imaging

Electrophysiological recordings in the on-cell and excised, inside-out mode were made with glass pipettes (WPI) that were that were generated with an electrode puller (Sutter Instruments) and fire polished with a microforge (Narishige) for a final resistance of 0.3-1 MOhm. Pipettes were filled with 4mM KCl, 136 mM NaCl, 1.8mM CaCl_2_, 1.0mM MgCl_2_, 5 mM HEPES, pH = 7.4 (the extracelluar solution) and the bath solution (intracellular solution in excised, inside-out mode) was 140 mM KCl, 5 mM HEPES, 5 mM MgATP_2_, pH = 7.4. Recordings were made with a EPC-10 patch clamp and PatchMaster software (HEKA). For PCF recordings, pipettes were imaged using a 40x water objective (Nikon) on a TE-2000U inverted microscope (Nikon) and illuminated with a SOLA light engine (Nikon). Pipette images were measured with a Prime 95B CCD camera, and with a custom CFP cube (436/20, dclp 455, D460lp) or YFP cube (500/20, Q515, HQ520lp) (Chroma). Spectra were measured using the appropriate cube and a SpectraPro 2150i spectrograph. Pipette images and spectra were obtained with NIS image acquisition and analysis software (Nikon).

### Measurement of FRET and analysis

FRET spectroscopy was performed using the custom CFP cube and YFP cube described above with spectra obtained with a spectrograph as in our previous studies (Codding & Trudeau, 2019; Gianulis et al, 2013). A value proportional to FRET efficiency, RatioA-RatioA_0_, was calculated from the spectra, as shown in our previous work (Codding & Trudeau, 2019; ^G^ianulis et al, 201^3^; Gustina & Trudeau, 2009; McNally et al, 2017; Trudeau & Zagotta, 2004) and as described (Selvin, 1995). For PCF mode, FRET was measured from *Xenopus* oocytes and for whole-cell mode, FRET was measured from HEK293 cells.

The relaxation of the current with repolarization (deactivation) was fit with a single exponential and GV curves were fit with a Boltzmann function. Curve fitting was performed using Igor (Wavemetrics) Statistical analysis was performed with GraphPad Prism where * denotes p ≤ 0.05 and ** denotes p ≤ 0.01 using a Students’ t-test. Error bars are the SE.

## RESULTS

To carry out these experiments we expressed hERG-Citrine, hERG ΔPAS-Citrine and hERG PAS - CFP *in trans* with hERG ΔPAS channels in *Xenopus* oocytes. All channels contained the S620T mutant, which reduced inactivation, in order to help obtain robust channel currents.

As a control, we recorded currents from patches expressing hERG-Citrine channels in the on-cell mode, immediately upon patch excision into the inside-out mode and then every 5 minutes over a total duration of 30 minutes (Fig. 2A-C). We found that the time constant of deactivation was similar over the duration of recordings (Fig. 2A-C, J). Likewise, as an additional control, we recorded hERG ΔPAS-Citrine channels in on-cell mode and then every 5 minutes over a total duration of 30 minutes. The time constant of deactivation was similar in on-cell and excised patches and was 5-fold faster than for hERG-Citrine channels, as anticipated, due to the lack of the regulatory PAS domain (Fig. 2D-F, K).

**Figure 2.**
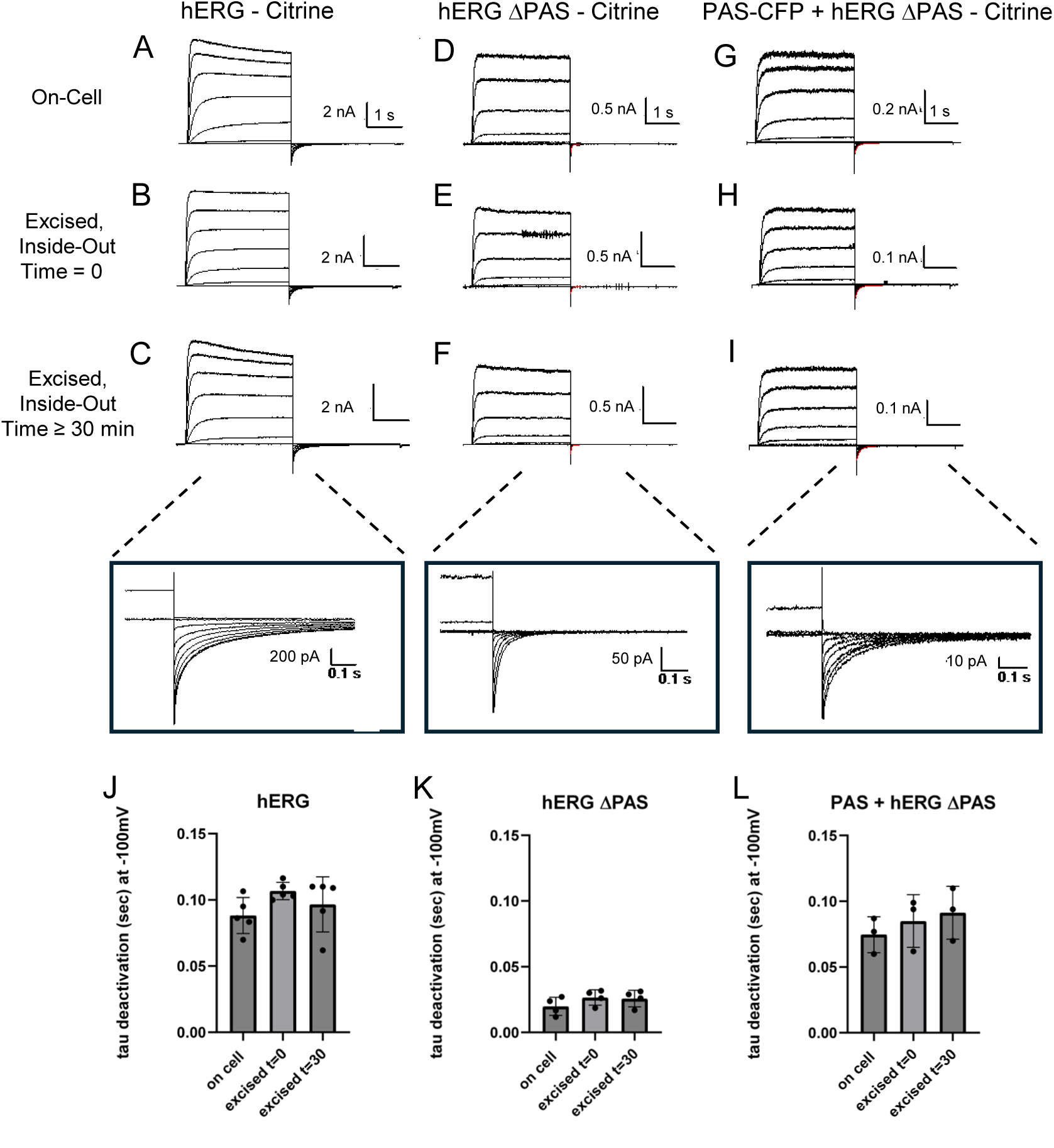
Deactivation is similar in on-cell and excised, inside-out patches for hERG and PAS + hERG. Δ**PAS channels.** Patch-clamp recordings in the on-cell configuration, immediately after patch excision and in the inside-out configuration 30 minutes following patch excision for (A-C) hERG-Citrine, (D-F) hERG ΔPAS-Citrine and (G-I) PAS-CFP+ hERG ΔPAS-Citrine. Insets show expanded view of tail currents after 30 minutes in excised, inside-out mode. Scatter plot of time constants of deactivation for (J) hERG-Citrine, (K) hERG ΔPAS Citrine and (L) PAS-CFP + hERG ΔPAS Citrine. Time constant of deactivation was derived from a fit to the inward tail currents at -100 mV. Error bars are SE and n ≥ 4 for each.

Next, we recorded currents from patches containing PAS-CFP + hERG ΔPAS-Citrine in on-cell mode, immediately upon patch excision, and then then at 5-minute intervals in the excised, inside-out mode for a duration of 30 minutes (Fig. 2G-I). We found that the time constant for deactivation was similar in on-cell and excised, inside-out configurations for the duration of the experiment (Fig. 2L). Moreover, deactivation for PAS-CFP + hERG ΔPAS-Citrine was similar to that for control hERG-Citrine channels and markedly slower than that for hERG ΔPAS-Citrine channels. Since the time constant for deactivation for PAS-CFP+ hERG ΔPAS remained the same over the course of the experiment, and was similar to that for control hERG-Citrine and slower than that for hERG ΔPAS channels we interpret this to mean that for PAS-CFP + hERG ΔPAS channels, the PAS domain remained attached to the channel and performed its regulatory function in the excised, inside-out patch-clamp mode in these experiments.

We also measured the conductance-voltage (GV) relationship in on-cell mode, immediately following patch excision, and 30 minutes after excision (Fig. 3). We did not detect a change in GV midpoints in different recording configurations in on-cell versus excised, inside-out configurations immediately after patch excision or after 30 minutes for hERG, hERG ΔPAS channels or PAS-CFP + hERG ΔPAS channels (Fig. 3A-C). But, compared to hERG channels, we detected an approximately 20-25 mV right-ward shift in the midpoint of the GV for hERG ΔPAS channels (Fig. 3A, B), which is consistent with the lack of the PAS domain, since a similar rightward shift was also detected in whole cell recordings of hERG ΔPAS channels compared to hERG channels (Gianulis et al, 2013; Gustina & Trudeau, 2011; Gustina & Trudeau, 2013; Spector et al, 1996; Wang et al, 1998). The midpoint of the GV for PAS-CFP + hERG ΔPAS channels was approximately -20 mV left-shifted compared to hERG ΔPAS channels and was similar to that of WT hERG channels (Fig. 3A-C) which also occurred in whole-cell measurements (Gianulis et al, 2013; Gustina & Trudeau, 2011; Gustina & Trudeau, 2013) and indicated that the PAS domain was regulating the channel, suggesting that the PAS domain was associated with hERG ΔPAS channels for the entirety of the experiment. The GV relationship results match with the slowed deactivation kinetics we observed in PAS-CFP + hERG ΔPAS Citrine channels (Fig. 2), and also indicated that the PAS domain remained associated with the channel during gating.

**Figure 3.**
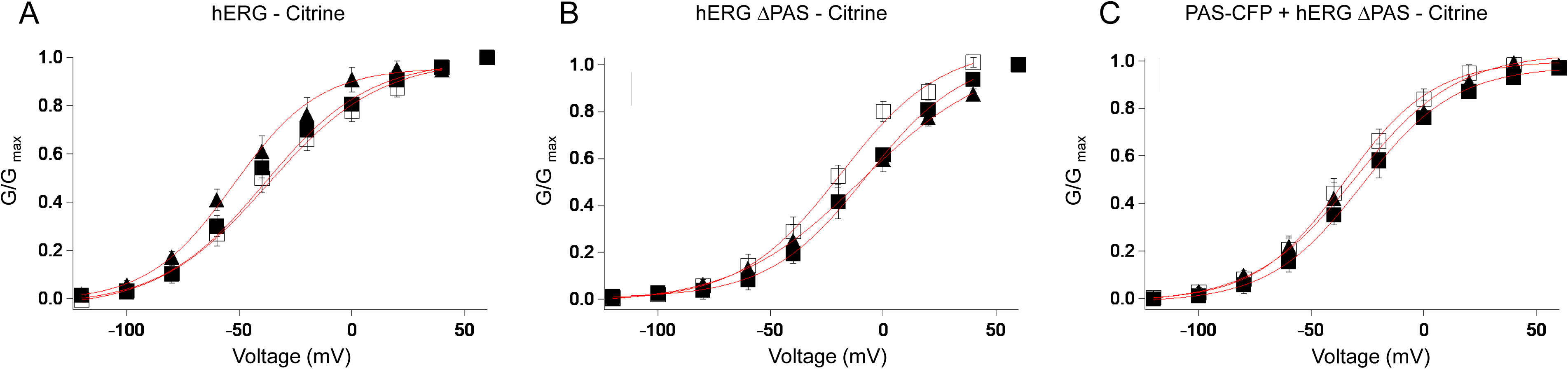
Steady-state activation is similar in on-cell and excised, inside-out patches for hERG and PAS + hERG ΔPAS channels. Plot of normalized conductance (G/Gmax) versus voltage (mV) in on-cell mode (solid square), immediately after patch excision (open square) and 30 minutes following patch excision (closed triangle) for A) hERG-Citrine, B) hERG ΔPAS Citrine and C) PAS-CFP + hERG Δ PAS Citrine. Error bars are SE and n≥4.

To obtain an independent measure for association of the PAS domain with the rest of the hERG channel we performed patch-clamp fluorometry (PCF) experiments. In PCF recordings, ionic currents and fluorescence are measured in the excised, inside-out patch-clamp configuration to simultaneously measure function and structure (Trudeau & Zagotta, 2004; Zheng & Zagotta, 2000; Zheng & Zagotta, 2003). As a positive control experiment, we performed PCF recordings from cells expressing hERG-Citrine or hERG-CFP channels (Fig. 4). In PCF images from cells expressing hERG-Citrine, we did not detect robust fluorescence with the CFP cube, but rather detected robust fluorescence with the YFP cube and we found that in cells expressing hERG-CFP that there was robust fluorescence detected using the CFP cube, but not with the YFP cube (Fig. 4A) and this was plotted as CFP fluorescence intensity versus Citrine fluorescence intensity (Fig. 4B) which indicated that CFP and Citrine fluorescence were distinguishable and separable. For hERG-Citrine channels we detected a linear relationship between ionic currents (Fig. 4C) and Citrine fluorescence intensity (Fig. 4 D) and for hERG-CFP channels we detected a linear relationship between ionic currents (Fig. 4C) and CFP fluorescence intensity (Fig. 4E), indicating that the fluorescence intensity was associated with channels in the membrane patch in these PCF experiments.

**Figure 4.**
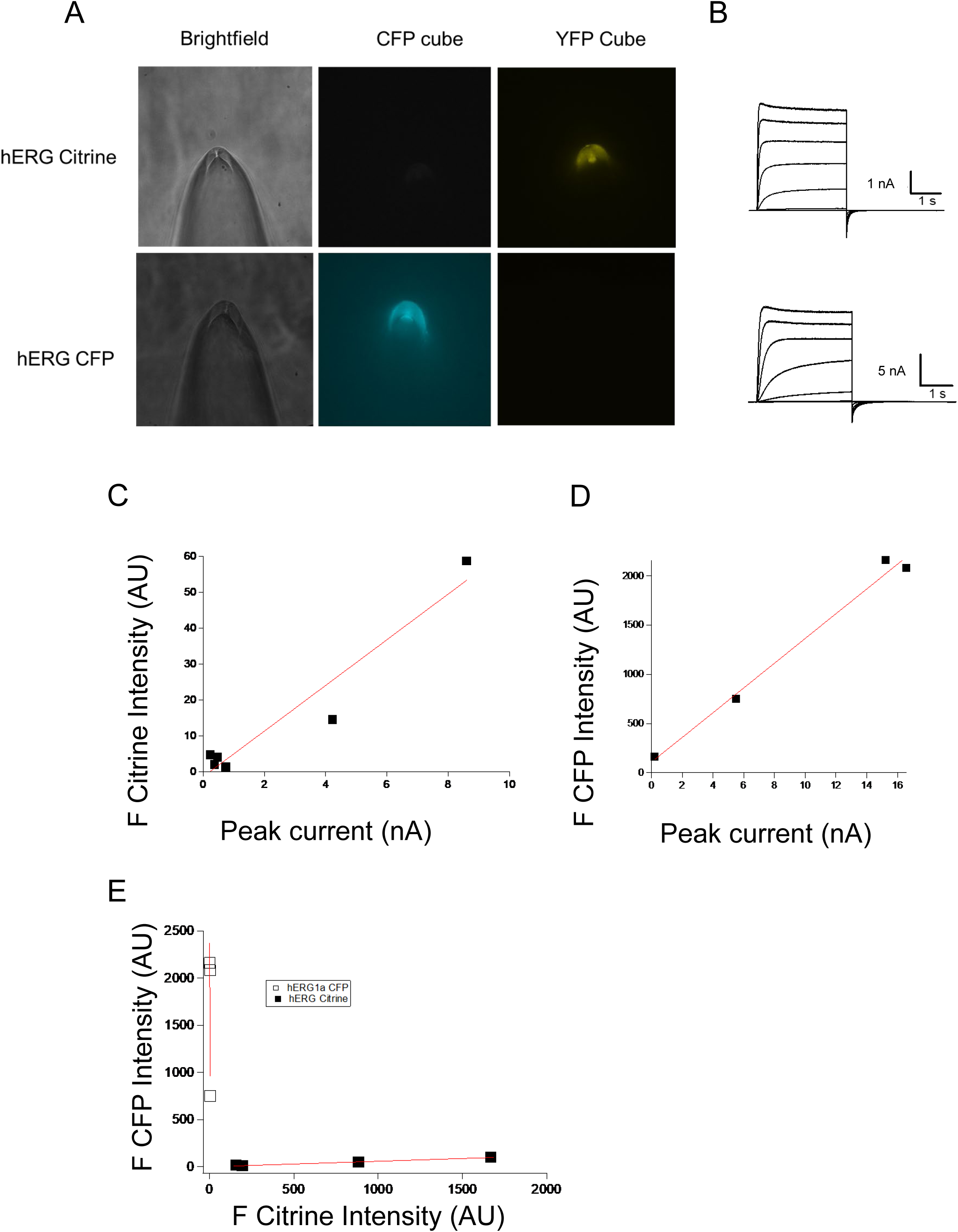
Patch-clamp fluorometry (PCF) recordings from hERG-CFP or hERG-Citrine channels. A) Brightfield images of patch pipette, fluorescence image with a CFP cube or fluorescence image with a YFP cube. B) Current recordings from hERG S620T-Citirne and hERG S620T-CFP in inside-out patch-clamp configuration. C) Plot of the CFP intensity versus Citrine intensity for hERG-CFP (open square) or hERG-Citrine (closed square). D) Plot of Citrine fluorescence intensity versus peak outward current at 60 mV for hERG-Citrine and E) Plot of CFP intensity versus peak outward current at 60 mV for hERG-CFP.

Next, as a negative control, we performed PCF recordings from oocytes expressing PAS-CFP and detected neither fluorescence associated with the membrane patch in either the CFP or Citrine cube nor ionic currents (Fig. 5A, B). We then performed PCF recordings from membrane patches with hERG ΔPAS-Citrine and did not detect a robust signal in the CFP cube, but detected robust fluorescence intensity in the YFP cube and robust ionic currents with rapid deactivation (Fig. 5A,B). In PCF recordings from patches expressing PAS-CFP + hERG ΔPAS-Citrine, we detected robust fluorescence intensity with both the CFP cube and the YFP cubes (Fig. 5A,B) and the currents measured simultaneously from the same patch had slower deactivation than that measured for hERG ΔPAS-Citrine channels (Fig. 5A, B). A plot of CFP intensity versus ionic currents for PAS-CFP and PAS-CFP + hERG ΔPAS -Citrine, indicated that robust fluorescence from PAS-CFP was only detected in the presence of hERG ΔPAS-Citrine, suggesting that PAS-CFP was not non-specifically associated with the membrane, but rather was directly and specifically associating with the hERG ΔPAS-Citrine channels in the patch (Fig. 6A). A plot of F_CFP_/F_Citrine_ intensity showed that only PAS-CFP + hERG ΔPAS-Citrine had robust intensity from both CFP and Citrine compared to negative controls with just PAS-CFP, which had no measurable signal, or hERG ΔPAS-Citrine which had robust Citrine fluorescence compared to CFP fluorescence (Fig. 6B). The F_CFP_/F_Citrine_ for control hERG-CFP (dashed cyan) and for hERG-Citrine (dashed black) were included for comparison.

**Figure 5.**
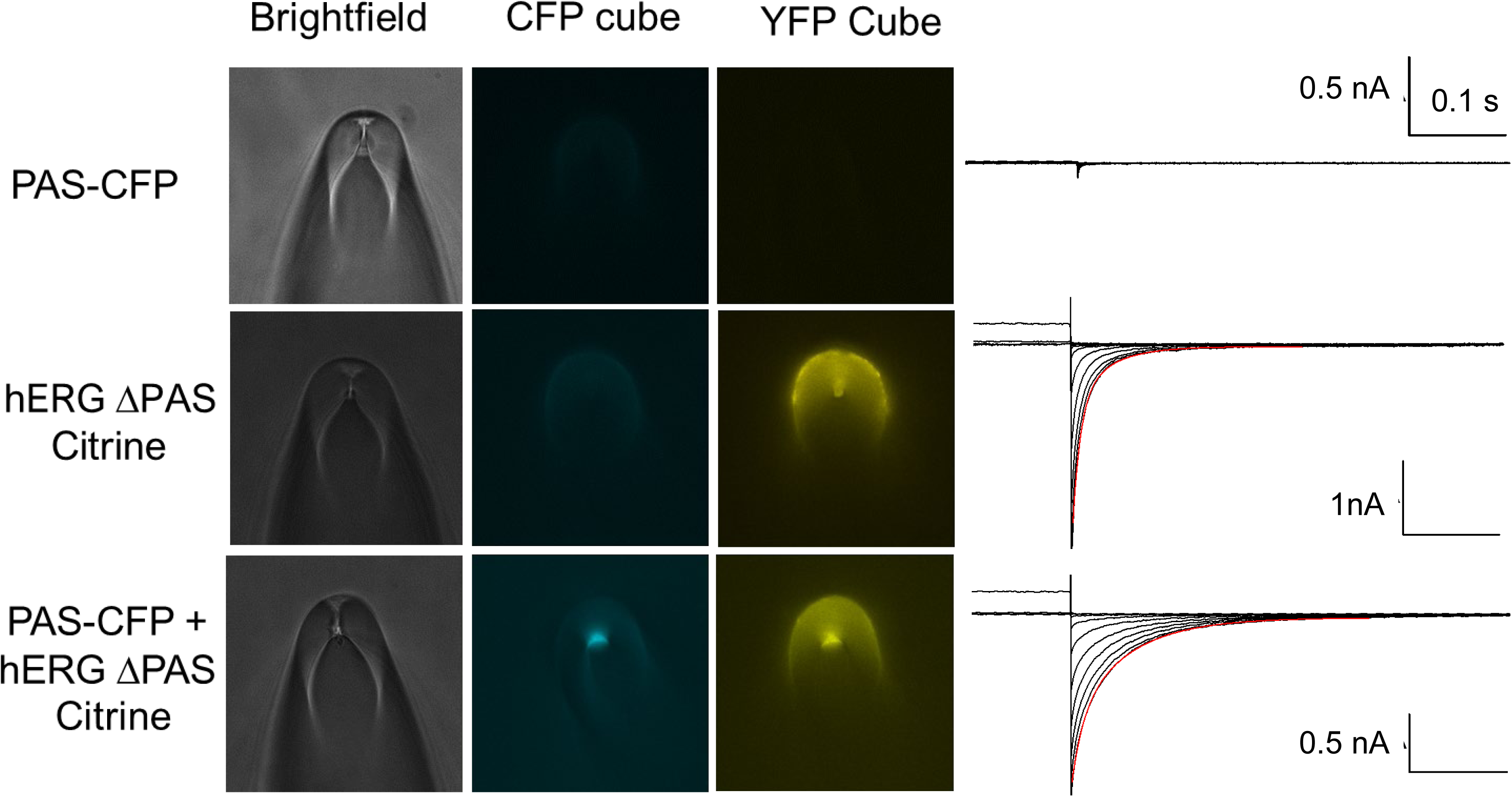
PCF recordings from PAS-CFP + hERG ΔPAS-Citrine channels. Images from patch pipette, membrane patches and corresponding currents from cells expressing PAS-CFP, hERG ΔPAS-Citrine and PAS-CFP + hERG ΔPAS-Citrine.

**Figure 6.**
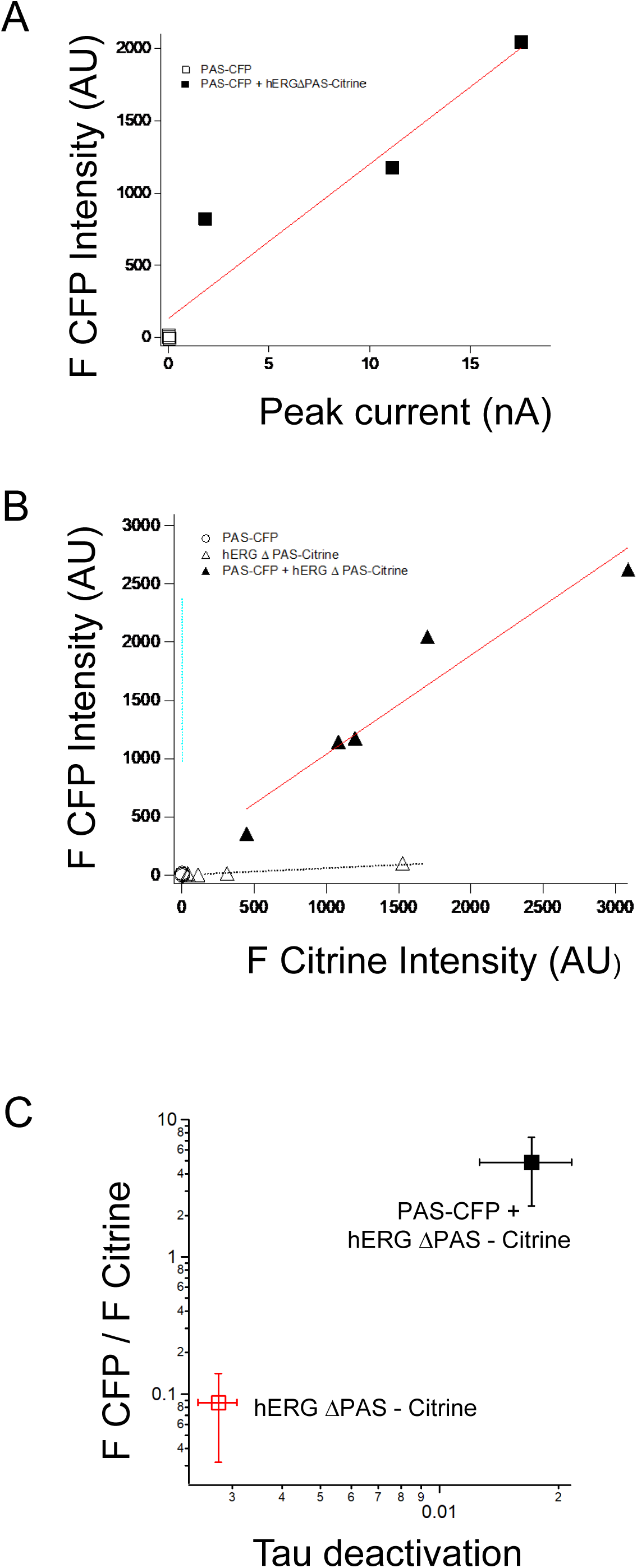
CFP fluorescence correlates with the time constant of deactivation for PAS-CFP + hERG ΔPAS-Citrine channels. Plots of A) CFP fluorescence intensity versus peak current at 60 mV for PAS-CFP (open square) and PAS-CFP + hERG ΔPAS-Citrine (closed square). B) Plot of CFP intensity (AU) versus Citrine intensity from PCF recordings for PAS-CFP (open circle), hERG ΔPAS-Citrine (open triangle), and PAS-CFP + hERG DPAS Citrine (closed triangle). Cyan dashed line is control hERG-CFP and dashed black line is control hERG-Citrine C) Plot of ratio of fluorescence from CFP and YFP cube (F _CFP_/F _Citrine_) versus time constant of deactivation for hERG Δ PAS-Citrine (red open square) and PAS-CFP + hERG ΔPAS-Citrine (closed black square). Error bars are mean ± SE.

A plot of the average ratio of F_CFP_/F_Citrine_ versus the time constant of channel deactivation showed that CFP fluorescence was correlated with slower deactivation (Fig. 6C). Compared to PCF recordings from hERG ΔPAS-Citrine channels, the increase in CFP fluorescence and slowing of deactivation in PCF recordings from PAS-CFP + hERG ΔPAS-Citrine channels both indicate that the PAS domain was present in these membrane patches.

To test more directly for an association between the PAS domain and the channel, we performed FRET experiments between PAS-CFP and hERG ΔPAS-Citrine (Fig. 7). We measured FRET from PCF recordings using ratiometric analysis of spectra, as in our previous studies (and see Methods). We first performed a control experiment by measuring the spectra from hERG ΔPAS-Citrine with the YFP cube (F_500_) and as a control for ‘bleed through’ from the CFP cube (F_436_). The control ratio of F_436_/F_500_ (Ratio A_0_) was 0.16 ± .01 which was consistent with previous studies (Fig. 7A). We next measured F_436_/F_500_ (Ratio A) from patches with PAS-CFP + hERG ΔPAS-Citrine, which was 0.37±.02, (Fig. 7B). We next calculated Ratio A-Ratio A_0_, which is proportional to FRET efficiency, and was 0.21 for PAS-CFP + hERG ΔPAS-Citrine in PCF mode, indicating robust FRET. As a positive control comparison, we determined that Ratio A-RatioA_0_ for PAS-CFP + hERG ΔPAS-Citrine in intact cells was similar to that in excised patches (Fig. 7C). These results suggest that PAS-CFP was associated with hERG ΔPAS-Citrine in excised patches, indicating that PAS remained attached to hERG channels.

**Figure 7.**
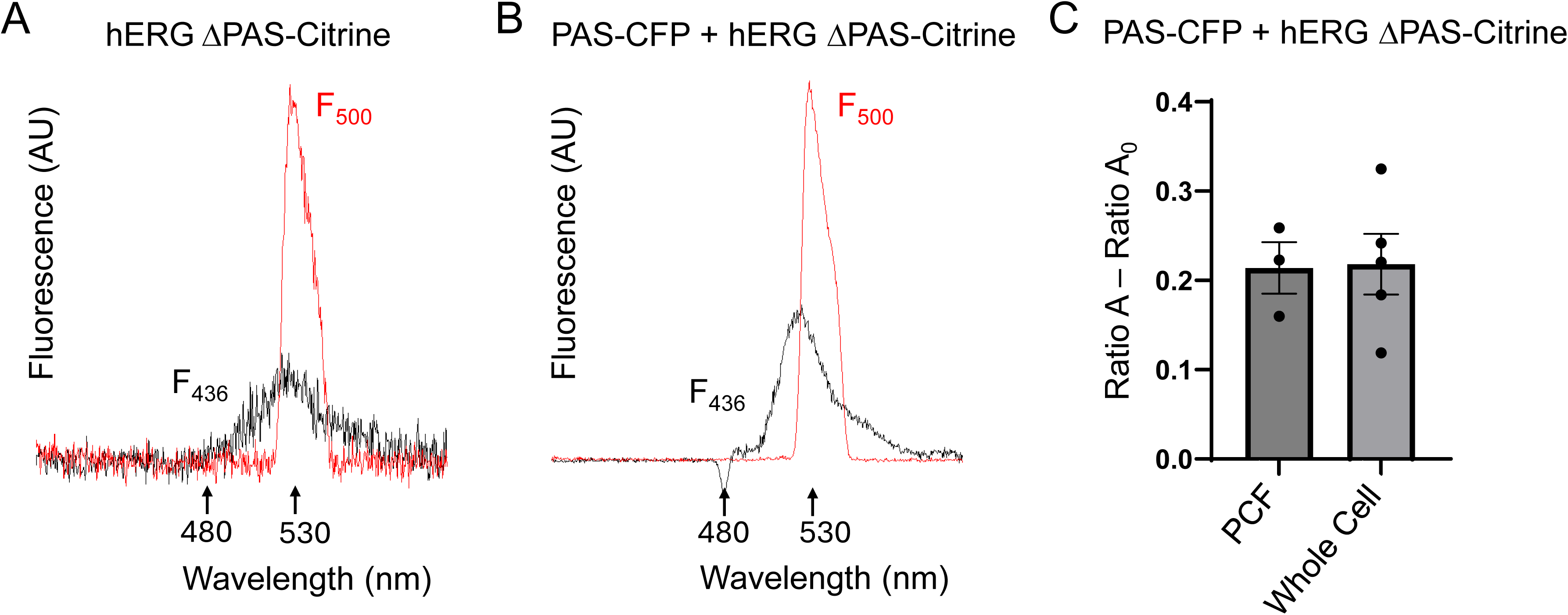
FRET in excised patches between the PAS-CFP and hERG DPAS Citrine channels. Plot of the spectra from the CFP/FRET cube (F_436_) or the YFP cube (F_500_) for A) control hERG ΔPAS-Citrine, B) PAS-CFP + hERG ΔPAS-Citrine channels, C) Plot Ratio A-RatioA_0_, for PAS-CFP + hERG ΔPAS-Citrine channels in excised, inside-out membrane patches (PCF mode) and in intact cells (whole cell mode).

**Figure 8.**
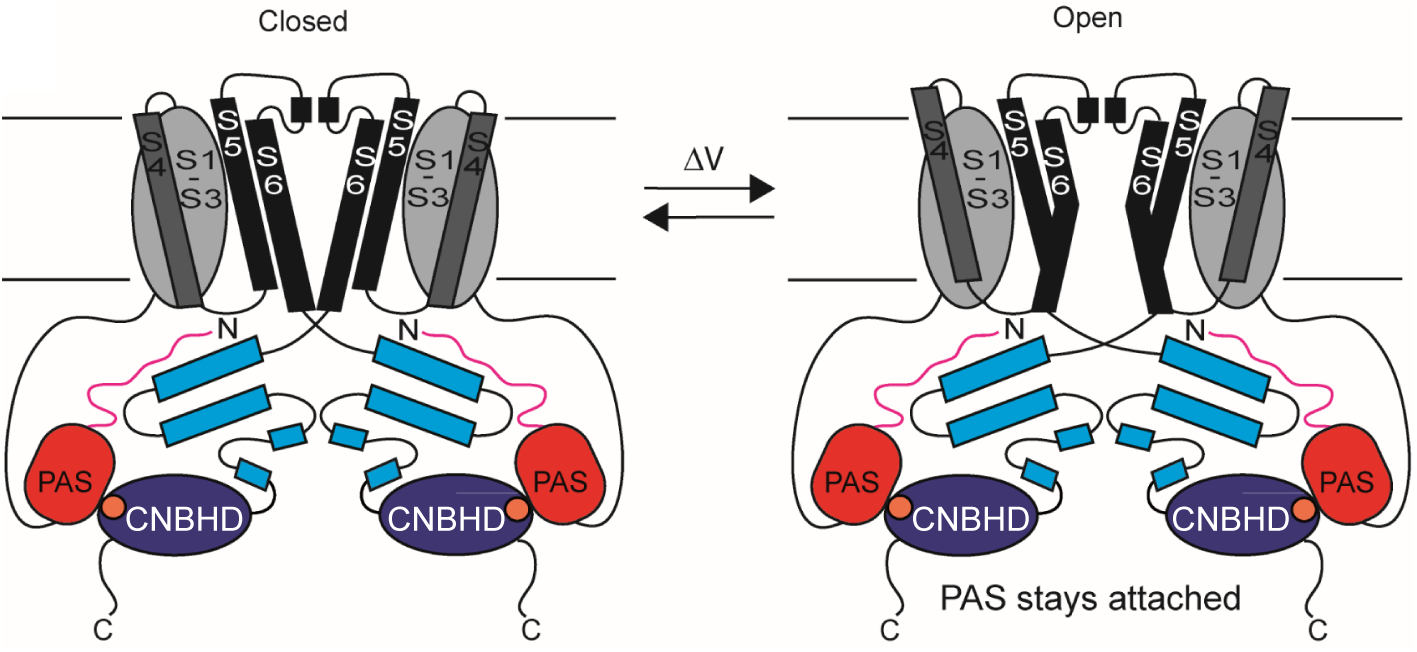
Schematic of mechanism for PAS domain regulation of gating in hERG. Schematic of Model I of gating in which PAS remains globally attached to the hERG channel during gating. Model II of gating in which PAS detaches during gating. Our results support Model I.

## DISCUSSION

Here we examined the mechanism for PAS domain regulation of hERG channel gating. We showed that PAS-CFP regulated hERG ΔPAS-Citrine channels in the excised, inside-out patch-clamp configuration. We found that for PAS-CFP + hERG ΔPAS-Citrine channels that deactivation was slow, like that for intact hERG-Citrine channels, and did not become fast, like that for hERG ΔPAS channels, after mechanical disruption of the membrane into the excised, inside-out mode. Deactivation remained slow for PAS-CFP + hERG ΔPAS Citrine channels despite long duration recording times in the excised, inside-out mode (30 minutes) or in response to voltage pulses to elicit different hERG conformational states. Likewise, the GV midpoint for hERG and PAS-CFP + hERG ΔPAS channels was similar in excised, inside-out mode, whereas that for the hERG ΔPAS channel was right-shifted by approximately 20-25 mV. Moreover, using PCF recordings, we detected robust CFP fluorescence and Citrine fluorescence in excised, inside-out patches containing PAS-CFP + hERG ΔPAS-Citrine, and robust CFP fluorescence was correlated with slow deactivation of currents. In contrast, in negative controls, we did not detect robust CFP fluorescence, but we detected robust Citrine fluorescence, and fast deactivation in hERG ΔPAS-Citrine channels. In a key control, we detected neither fluorescence nor currents from cells expressing PAS-CFP. Lastly, we detected robust FRET in PCF mode between PAS-CFP and hERG ΔPAS-Citrine.The PCF experiments provided an optical measurement (CFP fluorescence), in addition to a functional measurement (currents) which both indicated that the PAS was associated with hERG ΔPAS channels after patch excision.

The association of the PAS domain with the rest of the hERG channel in excised, inside-out patches suggests that PAS remained attached after mechanical disruption of the membrane. This suggests that PAS does not globally dissociate from the hERG ΔPAS channel, since if it did, it would be diluted in the infinite bath of the recording chamber and effectively lost and hERG deactivation would change from slow in PAS-CFP + hERG ΔPAS-Citrine patches to fast, as in hERG ΔPAS-Citrine patches, but we did not detect evidence that this occurred.

We extend our interpretation of these results to the mechanism for hERG gating and regulation by the PAS domain in intact, full-length channels. Since in PAS + hERG ΔPAS channels the PAS domain remained directly attached, this supports the mechanism for hERG gating in which PAS remains attached to the channel and does not detach during gating. This favors Model I for the mechanism of PAS gating (Fig.1).

Our conclusion that the PAS domain did not globally dissociate from the rest of the hERG channel during gating was consistent with other findings. In studies using transition metal FRET to probe PAS and CNBHD motions in ELK channels, which, like hERG, also have a PAS and CNBHD, it was found that the ELK channel PAS undergoes a small change in distance (a few Angstroms) relative to the CNBHD during gating (Dai et al, 2018). Moreover, using ultraviolet light to photo-inhibit PAS by crosslinking a photoactivatable, non-canonical amino acid incorporated into the hERG PAS domain there was a state-dependence to cross-link formation, which was also consistent with local motions of the PAS relative to the rest of the channel during gating (Codding & Trudeau, 2024). In this study, we did not detect a change in CFP fluorescence or FRET with application of voltage pulses, consistent with a previous study of FRET pairs placed into the N-and C-terminal regions of hERG channels in whole-cell mode (Kume et al, 2018). We do not interpret this to mean that the PAS domains do not move, but rather that FRET pairs based on large fluorescent CFP and Citrine proteins may not have the resolution to detect putative small motions associated with gating.

hERG channels appear to be a good example of modular functional domains in an ion channel. For instance, functional hERG channels can be formed from two channel halves that contain the N-terminus and VSD + the Pore module – COOH that are split at the S4-S5 linker and this is also the case for KCNH1 (EAG channels) (Lorinczi et al, 2015). Likewise, the PAS domain of hERG retains its regulatory function when not attached to the rest of the channel with a peptide bond, but instead exerts its function through a direct interaction with other intracellular domains, including the CNBHD (Codding & Trudeau, 2019; Gianulis et al, 2013; Gianulis & Trudeau, 2011; Gustina & Trudeau, 2009; Gustina & Trudeau, 2011; Gustina & Trudeau, 2012; Gustina & Trudeau, 2013; Morais Cabral et al, 1998; Trudeau et al, 2011). We propose that the excised, inside-out patch-clamp method with dual fluorescence (PCF) may be a useful approach to determining whether modular domains or accessory proteins remain attached to channels during gating or regulation. Thus, PCF with *in trans* expression of a regulatory domain (PAS) and a channel (hERG ΔPAS) may provide a more broad template for determining the stability and mechanism for direct intracellular domain interactions with ion channels.

## ACKNOWLEDGMENTS

Work was supported by NIH grants R01GM130701 and R01GM127523 to MCT.

